# Three-photon fluorescence microscopy with an axially elongated Bessel focus

**DOI:** 10.1101/240960

**Authors:** Cristina Rodríguez, Yajie Liang, Rongwen Lu, Na Ji

## Abstract

Volumetric imaging tools that are simple to adopt, flexible, and robust, are in high demand in the field of neuroscience, where the ability to image neurons and their networks with high spatiotemporal resolution is essential. Using an axially elongated focus approximating a Bessel beam, in combination with two-photon fluorescence microscopy, has proven successful at such an endeavor. Here we demonstrate three-photon fluorescence imaging with an axially extended Bessel focus. We use an axicon-based module which allowed for the generation of Bessel foci of varying numerical aperture and axial length, and apply this volumetric imaging tool to image mouse brain slices and for *in vivo* imaging of the mouse brain.

---

Two-photon fluorescence microscopy [1] has proven to be an invaluable tool for neuroscience, allowing for the study of neural structure and activity deep inside the living brain. More recently, three-photon fluorescence microscopy, in combination with long wavelength excitation, has been shown to push the depth limits of nonlinear imaging in scattering brain tissue [2, 3] and has allowed for transcutical imaging of the fly brain [4]. An attractive feature of these multiphoton imaging modalities relies in their optical sectioning capabilities. Since the excitation is localized to a small volume (on the order of a wavelength) around the focal plane, intrinsic 3D resolution is achieved. Neurons and their networks, however, can extend over hundreds of micrometers in three dimensions. In order to image a volume using a standard multiphoton fluorescence microscope, a stack of images at different depths must be obtained. Consequently, the volumetric acquisition speed is highly compromised, making it difficult to capture dynamic events, such as calcium transients [5], with subsecond temporal resolution. The need for faster volumetric imaging tools has motivated the design and implementation of a number of technologies [6]. Among them is the generation of an axially extended focus approximating a Bessel beam, by illuminating the back aperture of the excitation objective with an annulus of light [7]. With this approach, a projected view of the 3D volume is obtained, with a volume rate equivalent to the frame rate, without compromising the lateral resolution. Several approaches have been demonstrated for generating a Bessel beam under two-photon excitation. The most common approach involves the use of an axicon (conical lens) [8, 9, 10]. Other approaches employed a phase mask [11], or alternatively a spatial light modulator (SLM) [12], with the latter allowing for great flexibility in generating different types of Bessel foci with shaped axial intensity profiles.

In this paper, we demonstrate, for the first time to our knowledge, three-photon fluorescence imaging with an axially extended focus. We employ a flexible axicon-based Bessel module and demonstrate the generation of Bessel foci of different axial extents under three-photon excitation, and apply this volumetric imaging tool to image mouse brain slices and for *in vivo* imaging of the mouse brain.

A simplified diagram of our homebuilt three-photon microscope is shown in Fig. 1 **a**. The three-photon excitation source consists of a two-stage optical parametric amplifier (Opera-F, Coherent) providing a broad tuning range (650–900 nm and 1200–2500 nm) pumped by a 40 W diode-pumped femtosecond laser (Monaco, Coherent). For the experiments presented here, Opera-F was operated at 1300 nm, for which the average output power was ~1.5 W (1.5 μJ per pulse at 1 MHz repetition rate). To reduce the group delay dispersion at the sample plane, a homebuilt single-prim compressor was used [13]. The pulse duration at the focal plane of the objective was measured to be ~54 fs after compensation. A Pockel cell was used for controlling the power. A pair of galvanometers that were optically conjugate to each other and the back focal plane of a high-numerical aperture (NA) objective (Olympus XLPLN25XWMP2, NA 1.05, 25×) steered the excitation focus in the xy plane. The objective was mounted on a piezoelectric stage (PIFOC, Physik Instrumente) to translate the focus axially. The fluorescence signal is collected by the same objective and reflected from a dichroic beam splitter, spectrally filtered, and detected by a photomultiplier tube (Hamamatsu). Custom-written software was used for image acquisition.

**Fig. 1.**
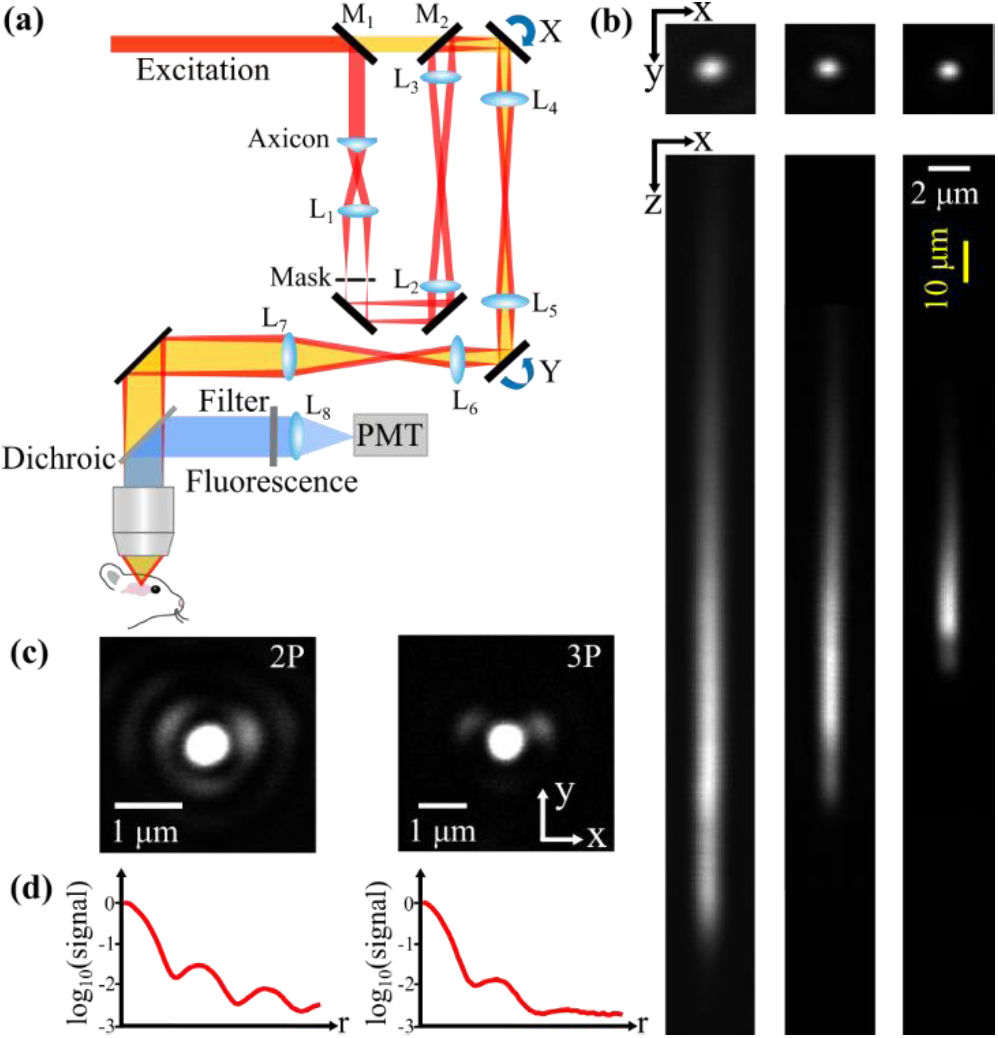
Three-photon fluorescence microscopy with a Bessel focus. **(a)** Simplified diagram of our homebuilt three-photon microscope with an axicon-based Bessel module. Flip mirrors M1 and M2 allow switching between Gaussian (yellow path) and Bessel (red path) imaging modalities. PMT, photomultiplier tube. **(b)** Three-photon images of a 1-μm-diameter red fluorescent bead with different Bessel foci: (left) NA = 0.48, focal length L_1_ = 100 mm, L_2_ offset = −10 mm; (center) NA = 0.6, focal length L_1_ = 125 mm, L_2_ offset = −10 mm; (right) NA = 0.72, focal length L_1_ = 150 mm, L_2_ offset = 0 mm Post-objective powers: 15.5, 4.9, and 3.2 mW, respectively. Note the different scales for the lateral and axial scale bars. **(c)** Images of a 0.2-μm-diameter red fluorescent bead taken under (left) two- and (right) three-photon excitation with a Bessel focus of 0.96 NA and axial FWHM of 10 μm (focal length of L_1_ = 200 mm, L_2_ offset = −10 mm). Contrast was enhanced 7× in both images. Post-objective powers: 10 and 16.7 mW, respectively. **(d)** Signal profiles (obtained from a radial average and plotted in logarithmic scale) of images in **(c)**. With three-photon excitation the side rings (typical of high-NA Bessel foci) are more effectively suppressed.

To generate a Bessel focus, we placed an axicon-based module before the scanning unit (Fig. 1 **a**). The module consists of an axicon (apex angle of 178°) followed by a lens (L_1_), which transforms the incident laser beam into an annulus of light. This annulus is imaged into the galvanometers (by the lens pair L_2_ and L_3_) and back focal plane of the objective, generating an axially elongated focus at the sample plane. An annular mask is placed at the back focal plane of lens L_1_ to block unwanted light arising from imperfections of the axicon. By changing the focal length of lens L_1_, the NA of the Bessel beam can be adjusted. Additionally, lens L_2_ can be translated along its optical axis, allowing for the adjustment of the effective NA as well as the axial length of the Bessel focus [10]. Here, moving lens L_2_ towards the mask (negative offset value) generates a longer Bessel focus; whereas moving lens L_2_ away from the mask (positive offset value) results in a shorter Bessel focus.

To illustrate the versatility of our Bessel module to generate Bessel foci of different NAs and consequently axial lengths, we imaged a 1-μm red fluorescent bead (Fluosphere; Invitrogen) using three different Bessel foci (Fig. 1 **b**) with an axial full width at half maximum (FWHM) ranging from 20 to 90 μm.

Previous observations showed that, for two-photon fluorescence microscopy, lower-NA (e.g., 0.4 or 0.6) Bessel foci produced better-quality images of brains *in vivo* [12]. For high-NA Bessel foci (e.g., 0.9), more energy is distributed in the side rings, causing a stronger background and image blur. Due to its higher nonlinearity, three-photon excitation should more effectively suppress the fluorescence signal generated by the side rings of a high-NA Bessel focus than two-photon excitation. Indeed, as illustrated in Fig. 1 **c**, the lateral (xy) images of a 0.2-μm red fluorescent bead (Fluosphere; Invitrogen) which were obtained using two- and three-photon excitation, respectively, showed substantially reduced side-ring signal via three-photon excitation for a 0.96-NA Bessel focus (Fig. 1 **d**). A titanium-sapphire laser (Chameleon Ultra II, Coherent) operating at 920 nm was used as the two-photon excitation source.

Having demonstrated the ability to generate Bessel foci under three-photon excitation, we applied this technique to image mouse brain slices (Thy1-GFP line M). Using a Bessel focus with axial FWHM of 36 μm and 0.6 NA, we were able to image neural structures extending ~50 µm in depth in a single frame (Fig. 2). In Fig. 2 **b**, the Bessel image captured the basal dendrites surrounding the neuronal cell body; in Fig. 2 **d**, the Bessel image resolved fine dendrites and dendritic spines over a similar volume. In comparison, a Gaussian focus, generated by overfilling the back aperture of the 1.05 NA microscope objective with 1300 nm excitation, has an axial FWHM of ~1.8 μm under three-photon excitation. In order to capture all the structures in the same volume as that obtained with a Bessel focus (Fig. 2 **b,d**), we acquired ~50-μm thick image stacks made of 26 frames (Fig. 2 **a,c**, color-coded by depth), with 2-μm axial step size. Comparing the Gaussian and Bessel images indicates that finer structures visible in the Gaussian stacks were clearly resolved using the Bessel imaging modality (arrowheads in Fig. 2 **c,d**). Therefore, for three-photon fluorescence microscopy, scanning a Bessel focus captures all features of interest in a volume in a single frame, without compromising the lateral resolution.

**Fig. 2.**
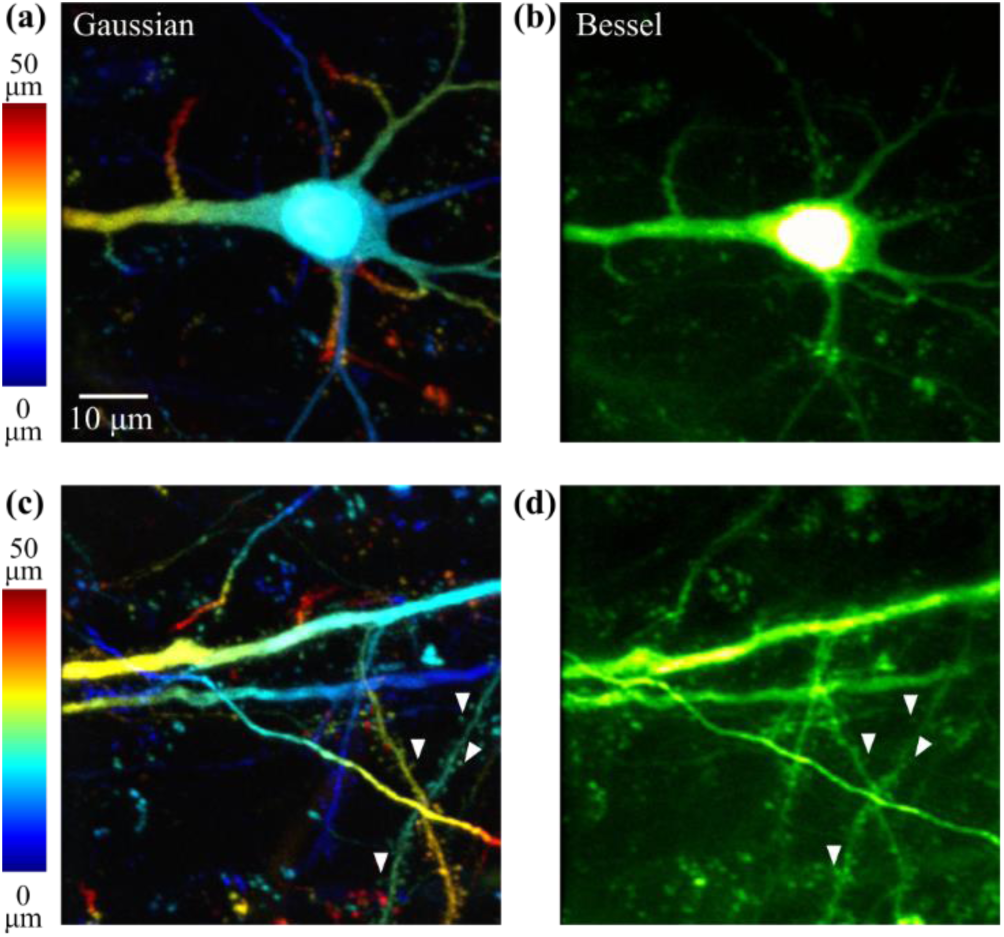
Three-photon microscopy images of mouse (Thy1-GFP line M) brain slices imaged with a Gaussian or Bessel focus. **(a,c)** Maximum intensity projections of a 50-μm thick image stack obtained with a Gaussian focus, with a 2-μm axial step size, color-coded by depth. Post-objective power: 2.8 mW. **(b,d)** Images of the same volumes as **(a,c)**, respectively, obtained with a Bessel focus of NA 0.6 and axial FWHM of 36 μm. Post-objective powers: 25 and 38 mW, respectively.

Imaging biological structures *in vivo* is one of the most valuable applications of optical microscopy, allowing for the study of biological systems in a relatively undisturbed and more physiological state. Figure 3 shows representative images taken during *in vivo* three-photon imaging of the mouse brain (Thy1-GFP line M). Using a Bessel focus having an axial FWHM of 38 μm, we were able to probe neural structures extending over 60 μm *in vivo* in a single frame, while maintaining the lateral resolution to resolve synaptic structures such as dendritic spines (white arrowheads) and axonal boutons (yellow arrowheads). Similar to our observations in brain slices, all the fine structures observable in the image stacks obtained with a Gaussian focus were clearly identifiable using the Bessel imaging modality. It is important to note that glass cranial windows, used for providing optical access into the mouse brain, introduce aberrations. If not compensated for, such aberrations have a detrimental effect on the image quality. Here, we used the correction collar of the objective to compensate for the spherical aberration introduced by the glass window. Furthermore, to avoid the additional aberration modes that arise from a tilted cranial window (e.g., coma, astigmatism) [14], we carefully positioned the mouse using the third-harmonic signal from the glass window interfaces to check and correct for any tilt. For application in deeper depths, we will utilize adaptive optics to correct for the sample-induced aberrations [4, 15, 16].

**Fig. 3.**
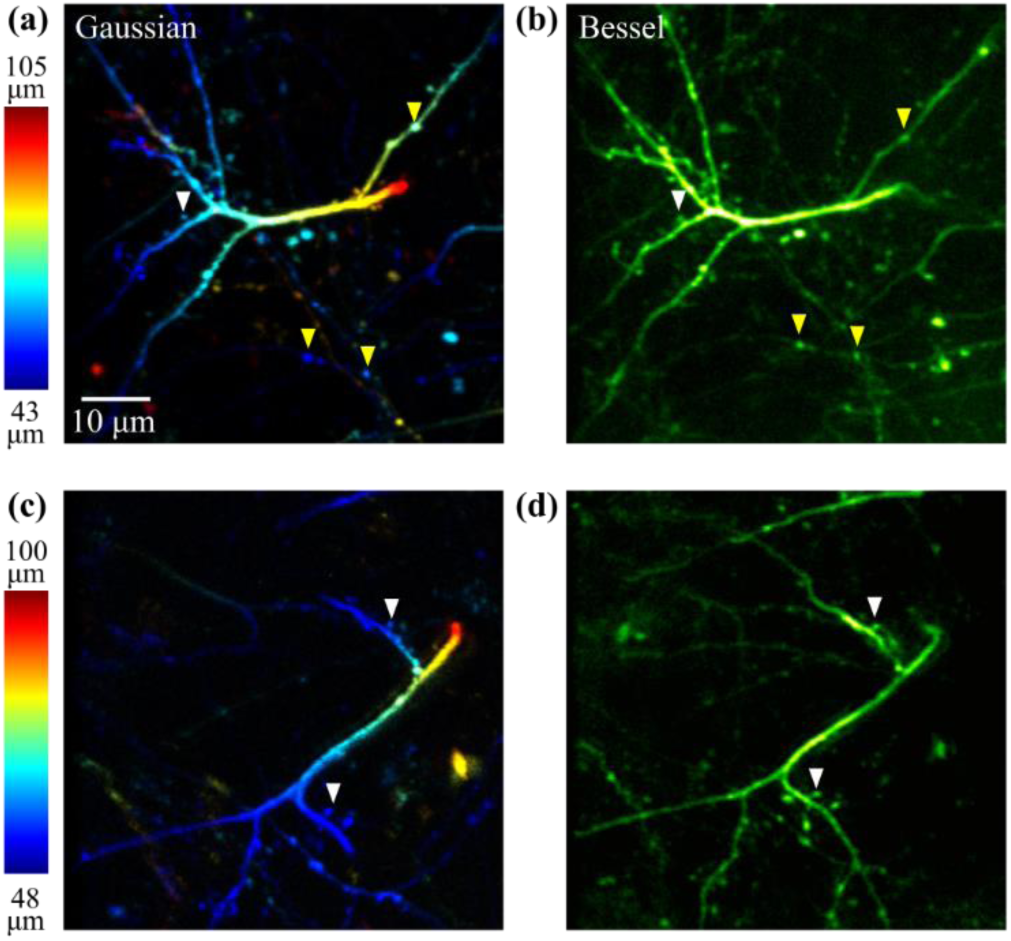
*In vivo* three-photon microscopy of a mouse (Thy1-GFP line M) brain with a Bessel focus. Maximum intensity projection of a **(a)** 62-μm and **(c)** 52-μm thick image stack obtained with a Gaussian focus, with 2-μm axial step size, color-coded by depth, ~80 μm below dura. Post-objective powers: 3 and 4 mW, respectively. **(b,d)** Image of the same volume as **(a,c)**, respectively, obtained with a Bessel focus of NA 0.6 and axial FWHM of 38 μm. White arrowheads label dendritic spines, and yellow arrowheads label axonal boutons. Post-objective powers: 30 and 40 mW, respectively.

We have demonstrated three-photon fluorescence imaging with an axially extended focus. An axicon-based Bessel module was incorporated into a homebuilt three-photon microscope, which allowed for the generation of Bessel foci of varying NA and axial extents. When compared to two-photon excitation, three-photon excitation was shown to more effectively suppress the signal associated with side rings of high-NA Bessel foci. Both in fixed slices and *in vivo*, three-photon excitation fluorescence microscopy using a Bessel focus allowed us to capture volumetric data at high throughput while maintaining synaptic resolution. Another advantage of using a Bessel focus for *in vivo* brain imaging is the robustness of this volumetric imaging tool against axial motion. With an axially extended focus as the excitation, structures of interest can remain within the imaged volume even in the presence of small axial displacements of the sample, which are otherwise not straightforward to correct for with a Gaussian focus [17].

## Acknowledgment.

We thank the Ji lab for helpful discussions, and Erminia Fardone for cutting the brain slices.

